# Ectopic expression of pericentric HSATII RNA results in nuclear RNA accumulation, MeCP2 recruitment, and cell division defects

**DOI:** 10.1101/2020.04.30.064329

**Authors:** Catherine C. Landers, Christina A. Rabeler, Emily K. Ferrari, Lia R. D’Alessandro, Diana D. Kang, Jessica Malisa, Safia M. Bashir, Dawn M. Carone

## Abstract

Within the pericentric regions of human chromosomes reside large arrays of tandemly repeated satellite sequences. Expression of the human pericentric satellite HSATII is prevented by extensive heterochromatin silencing in normal cells, yet in many cancer cells, HSATII RNA is aberrantly expressed and accumulates in large nuclear foci *in cis*. Expression and aggregation of HSATII RNA in cancer cells is concomitant with recruitment of key chromatin regulatory proteins including methyl-CpG binding protein 2 (MeCP2). While HSATII expression has been observed in a wide variety of cancer cell lines and tissues, the effect of its expression is unknown. We tested the effect of stable expression of HSATII RNA within cells that do not normally express HSATII. Ectopic HSATII expression in HeLa and primary fibroblast cells leads to focal accumulation of HSATII RNA *in cis* and triggers the accumulation of MeCP2 onto nuclear HSATII RNA bodies. Further, long-term expression of HSATII RNA leads to cell division defects including lagging chromosomes, chromatin bridges, and other chromatin defects. Thus, expression of HSATII RNA in normal cells phenocopies its nuclear accumulation in cancer cells and allows for the characterization of the cellular events triggered by aberrant expression of pericentric satellite RNA.

## Introduction

Nearly 50% of the human genome consists of repetitive DNA sequence elements. These include transposable element-based repeats such as Long and Short Interspersed Nucleotide Elements (LINES and SINES) and simple sequence repeats (tandem DNA) such as ribosomal DNAs and satellite sequences (Richard et al., 2008). Distinct classes of repeat elements are uniquely localized within the genome. While LINE and SINE elements are heterogeneously interspersed genome-wide, tandemly repeated elements are more positionally defined. Satellite DNAs, defined by tandemly repeating units of DNA, localize primarily to centric or pericentric regions and to telomeres of all chromosomes, and are classified based on their sequence and length of repeating unit. The major classes of human satellite DNA are: 1) alphoid (alpha satellite; *α*-Sat) DNA resident at all centromeres; 2) telomeric repeats; 3) beta satellite DNA (*β*-Sat) on acrocentric chromosomes; and 4) satellites HSATI, HSATII, and HSATIII found at the pericentric regions of a subset of chromosomes.

Large blocks of HSATII DNA are located on 11 human chromosomes, with the largest arrays residing within the pericentromeres of human chromosomes 1 and 16. Satellite DNA blocks may span several megabases and consist of tandemly repeated elements ranging from five to several hundred nucleotides (Richard et al., 2008). Length and sequence composition of satellites, including HSATII, varies in the human population and may be used to study human genetic variation (Miga, 2019); however, the full extent and diversity of HSATII sequences is currently unknown due to its location within large sequence assembly gaps in the human genome (Altemose et al., 2014; Eichler et al., 2004). HSATII is roughly defined as a ∼26 bp repeat, but exhibits significant heterogeneity in sequence and has been poorly studied, despite its prominence in the human karyotype (Tagarro et al., 1994) and abundance in the human genome. A computational effort to generate a satellite reference database (Altemose et al., 2014) successfully demonstrates the ability to cluster HSAT sequences into subfamilies, which are likely to reside on individual chromosomes, but a comprehensive HSATII map has yet to be completed. In contrast, significant progress has been made in mapping alpha satellite arrays, consisting of 171-bp monomer repeats, particularly on the X (Miga et al., 2019) and Y (Jain et al., 2018) chromosomes, to generate full telomere-to-telomere linear chromosome sequence.

Due to their localization near centromeres, satellite sequences are likely to be subject to strict regulation to maintain both genetic and epigenetic stability. While expression of pericentric satellites has been observed in many species, including yeast, plants and mammalian cells, it is clear that when satellites are expressed, their expression is highly regulated (reviewed in (Hall et al., 2012; Perea-Resa and Blower, 2018; Smurova and De Wulf, 2018)). The increased presence of satellite RNA has recently emerged as an indicator of instability, as demonstrated by satellite overexpression in cancer cells and tumor tissues (Hall et al., 2017; Ting et al., 2011; Zhu et al., 2011). Whereas *α*-sat is expressed at low levels in normal cells (Hall et al., 2017; McNulty et al., 2017) and increases expression in cancer, HSATII expression is restricted to cancer cells, and thus the presence of HSATII RNA is a potential biomarker of cancer (Hall et al., 2017; Ting et al., 2011). Overexpression of chromosome-specific pericentric HSATII loci (e.g. Chr 7) occurs within nuclei in which other HSATII chromosomal locations (e.g. Chr 1) are not expressed, due to their accumulation of repressive polycomb proteins, thus indicating that HSATII expression is regulated in a locus-specific manner (Hall et al., 2017). Accumulation of HSATII RNA occurs *in cis*, with the RNA accumulating in cancer-associated satellite transcript (CAST) bodies, adjacent to the HSATII locus from which it is transcribed. Within a given tumor or cancer cell line, expression of HSATII maintains this locus-specific expression pattern in most cells (Hall et al., 2017).

HSATII RNA within CAST bodies recruit and directly bind MeCP2 and its protein-binding partner, Sin3A. (Hall et al., 2017). MeCP2 is canonically classified as a DNA methyl-binding repressor protein (Hite et al., 2009), however, more recent evidence indicates that MeCP2 also contains an intrinsically disordered domain, which has the capacity to bind RNA (Castello et al., 2016), and can function as a transcriptional activator (Hite et al., 2009). MeCP2 is mutated in Rett Syndrome, where it is known to bind transcripts in the brain and affect alternative splicing, thus suggesting MeCP2 has multifunctional roles in gene regulation and splicing (Young et al., 2005). MeCP2 is recruited to HSATII RNA accumulations and may be sequestered in CAST bodies in cancer cells, but the dynamics, consequences, and direct role of HSATII RNA in this recruitment are currently unknown.

Inappropriate expression of satellite RNAs can be an indicator of heterochromatic instability, which may be increasingly common in cancers, and has wide-ranging implications (Carone and Lawrence, 2013). Given that maintenance of centric/pericentric heterochromatin is key to normal chromosome segregation, structural changes to the underlying chromatin may result in functional changes, including satellite expression and aberrant cell division. Supporting this, prior studies analyzing the effect of satellite expression have demonstrated that forced expression of *α*-sat transcripts leads to cell division defects, chromosomal instability and aneuploidy in normal cells (Chan et al., 2017; Ichida et al., 2018; Zhu et al., 2018). Paradoxically, some *α*-sat expression is observed in normal cells (Hall et al., 2017) and low levels of expression are thought to be necessary for centromere function and constitutive heterochromatin maintenance (Johnson et al., 2017; McNulty et al., 2017). Thus, the functional role of satellites has been an enigma, despite their prevalence and structural conservation within pericentric regions. This conserved, tandemly repetitive structure of centric and pericentric satellites is in direct juxtaposition to the demonstrated lack of sequence conservation between species (Henikoff et al., 2001) and the discovery of functional neocentromeres lacking satellite sequences (Voullaire et al., 1993). Pericentric satellites, in particular, have little known function despite their abundance within the pericentric sequence landscape. It has become clear that the mechanisms governing transcriptional silencing of pericentric satellites are complex, with many histone modifications contributing to maintaining repression. Thus, a ubiquitous role for pericentric repeats, the mechanisms governing their regulation, and the satellite sequences that comprise them, has remained poorly understood. Given that pericentric satellite expression is misregulated in a wide range of cancer cell lines and tissues, and the chromatin structure of pericentric satellites is compromised in both cancer and senescent cells (Brückmann et al., 2018; Hédouin et al., 2017; Slee et al., 2012; Swanson et al., 2013; Tasselli et al., 2016), the direct effect of aberrant pericentric satellite expression warrants investigation.

Here, we established cell lines stably expressing HSATII RNA and *α*-sat RNA in order to test the long-term effects of expression of these distinct satellite transcripts. We demonstrate that ectopically expressed HSATII RNA accumulates in nuclear bodies reminiscent of CAST bodies in both HeLa and primary fibroblast cells. The focal accumulation of HSATII transcripts is unique to HSATII, as stable ectopic expression of *α*-sat RNA does not result in focal RNA accumulation in the nucleus. Further, HSATII RNA accumulates *in cis*, immediately adjacent to the genomic integration site, and recruits MeCP2 to these nuclear foci. Expression of both *α*-sat and HSATII transcripts leads to cell division defects including chromatin bridges, blebbing and micronuclei formation, thus indicating that expression of both centromeric and pericentric satellite sequences has the potential to generate chromosomal instability and aberrant cell division. The study of the effect of HSATII expression represents a unique opportunity to shed light on the function of pericentric satellite sequences more generally, in addition to testing the functional consequences of expression of pericentric satellite sequences in cancer.

## Results

### Establishment of a cell culture system to study the effect of HSATII RNA expression

Expression of HSATII RNA in cancer cells has previously been observed in both cancer cells grown in culture and from tissue biopsies (Hall et al., 2017; Ting et al., 2011). While there are distinct loci that harbor HSATII DNA sequences within the pericentric regions of roughly 11 human chromosomes, only a subset of these HSATII sequences are expressed, and the HSATII RNA expressed from these loci remain *in cis* in CAST bodies. HSATII sequences resident on human chromosomes 7 and 10 are among those sequences displaying a preference for expression in a variety of different cancer cell lines and tissues (Hall et al., 2017). In order to study the direct effect of expression of HSATII RNA, we developed a cell culture model to stably express HSATII sequence derived from Chr 7 in cell lines that do not endogenously express HSATII. To further examine the effect of HSATII expression irrespective of its location of expression, stable cell lines were created in which the Chr 7 HSATII-expression construct had been randomly integrated into the genome.

An HSATII cDNA sequence derived from Chr 7 was cloned into a plasmid designed for mammalian expression and stable integration, containing a CMV promoter and a neomycin selectable marker **(Figure 1A)**. HeLa cells, while a cancer cell line, do not endogenously express HSATII RNA at significant levels (Hall et al., 2017), thus initial transfection experiments were conducted in HeLa cells due to their ease of transfection by lipid-mediated transfection. To test for HSATII-specific effects, control vectors containing an *α*-sat sequence derived from Chr 4 and no insert (empty vector) were also used concurrently to transfect HeLa cells. Cells were assayed for transient satellite expression 24 hours after transfection by RNA FISH and RT-qPCR. 24 hours after transfection, approximately 20% of HSATII-transfected HeLa cell nuclei displayed nuclear accumulations of HSATII RNA compared to less than 5% of *α*-sat and empty-vector control transfected cells **(Figure 1B, D)**. Nucleoplasmic and cytoplasmic diffuse expression was also observed at this early timepoint, likely due to high levels of expression driven by the CMV promoter. Cells transfected with *α*-sat displayed a similar level of expression, with roughly 23% of cells displaying *α*-sat RNA by RNA FISH **(Figure 1C, E)**. However, a striking difference was observed in the distribution of HSATII and *α*-sat RNA in the nucleus. Distinct focal accumulations of HSATII RNA (2-3 per nucleus on average) were observed **(Figure 1B)**, while *α*-sat RNA appeared as a diffuse, primarily nuclear RNA signal **(Figure 1C)**. Expression of HSATII or *α*-sat was dependent on transfection with the respective insert-containing vector, thus demonstrating construct delivery specificity. Further, the percentage of cells expressing the desired sequence insert was significantly different from controls (empty vector) **(Figure 1F)**. RT-qPCR also confirmed high levels of HSATII expression in HSATII-transfected cell lines compared to alpha-sat transfected and controls **(Figure 1G)**. Since RT-qPCR was performed from total cellular RNA, results here cannot distinguish between nuclear RNA accumulations and diffuse RNA (nuclear or cytoplasmic), thus the greater than 8-fold increase in HSATII expression in HSATII-transfected cells likely illustrates the total amount of HSATII overexpression compared to *α*-sat and control cells (**Figure 1G)**.

**Figure 1.**
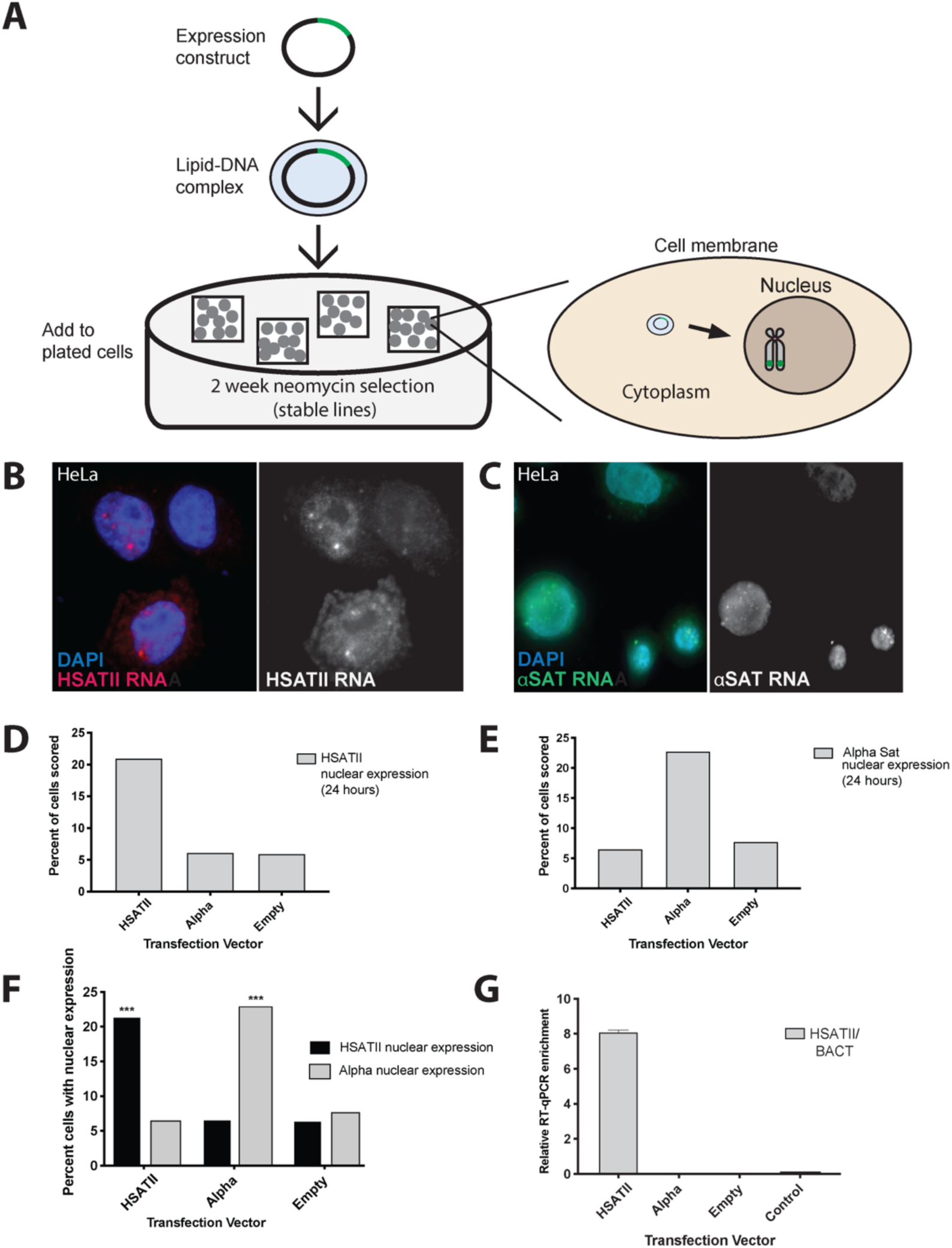
Transient transfection of satellite expression results in nuclear satellite RNA accumulation. **A)** Transfection scheme for transient and stable integration expression. A plasmid harboring HSATII, *α*-sat (*α*sat), or no insert (empty vector) is introduced to cultured HeLa or Tig-1 primary fibroblast cells via lipid-mediated transfection and cells are then fixed on coverslips or harvested for RNA isolation. Stable cell lines are further selected with neomycin (G418) for 2 weeks prior to fixation or harvesting. 24 hours after transfection, nuclei are scored for expression of **B)** HSATII and **C)** *α*-sat RNA signal by RNA FISH. Percent of cells (out of 500 nuclei) with **D)** HSATII RNA nuclear expression and **E)** *α*-sat nuclear expression. **F)** Nuclear RNA signal detected by RNA FISH is dependent on the sequence harbored within the transfected construct. Asterisks denote significant differences by unpaired t-test, p<.001. **G)** qRT-PCR of HSATII RNA 24 hours after transfection. Expression values shown relative to housekeeping gene expression (ß-actin) for each transfection condition.

While HeLa represent an easily transfected cell line that does not express HSATII RNA, they are cancer cells that have been maintained in culture since the 1950s, thus are not representative of “normal” cells. Following successful demonstration of transient HSATII and *α*-sat expression in HeLa cells, stably transfected cell lines were generated for both HeLa and a primary (non-transformed) human fibroblast cell line to ensure the results observed were not simply an effect of expressing satellite RNA in cancer cells, in which the nuclear and chromatin environment can be drastically different from normal cells (Carone and Lawrence, 2013; Zink et al., 2019). Tig-1 cells are female primary fibroblasts maintained at low passage number that retain an inactive X chromosome and a stable karyotype (2n= 46) (Ohashi et al., 1980), and can be transfected using lipid-mediated transfection. A lipid-mediated transfection protocol was optimized for successful transfection of Tig-1 primary fibroblasts, to ensure high transfection efficiency in addition to high cell viability (**Supplemental Figure 1**).

Following two weeks of neomycin selection, mock transfected and control HeLa and Tig-1 cultures displayed complete cell death, while a subset of viable cells (ranging from 10-15%) harboring HSATII, *α*-sat and empty vector integrants were selected and further expanded for a period of several weeks. Cells from stably transfected cell lines were fixed for DNA/RNA FISH and pelleted and harvested for RNA extraction at weekly intervals to assay satellite expression (**Figure 1A**). To test for genomic integration of the expression vector, DNA hybridization was performed using a biotin labeled probe complementary to the plasmid (pTargeT) backbone, which indicated that the majority of cells with detectable signal had one or two sites of integration (**Supplemental Figure 2**). Analysis of chromosome spreads harvested from transfected cell lines confirmed random integration of the HSATII expression vector, with the majority of spreads with hybridization signal harboring one or two integration sites, which were randomly distributed on chromosomes (**Supplemental Figure 3**). Three weeks following transfection, HeLa cells retained the same pattern of satellite expression, with *α*-sat RNA being diffuse and nuclear **(Figure 2A)**. In contrast, ∼5% of nuclei in HSATII-transfected cells displayed nuclear accumulations of HSATII RNA. This pattern was similarly observed in Tig-1 primary fibroblasts, with 7% of HSATII transfected cells harboring nuclear accumulations of HSATII RNA (**Figure 2E, I**) and *α*-sat RNA displaying a diffuse nuclear signal by RNA FISH (**Figure 2D**). RT-qPCR further confirmed higher levels of HSATII RNA in HSATII-transfected HeLa (**Figure 2H**) and Tig-1 (**Figure 2J**) cells. Of note, while transiently transfected cell lines displayed some cytoplasmic satellite RNA, stably transfected cell lines had very little cytoplasmic satellite RNA (*α*-sat or HSATII), suggesting that the vast majority of satellite RNA transcribed ectopically in these cell lines is retained in the nucleus. Taken together, these results indicate that stably expressing HeLa and primary fibroblast cell lines can be used to examine the effect of satellite RNA (HSATII and *α*-sat) expression in cells that do not normally express HSATII. Further, results from both transient and stably transfected cell lines indicate that HSATII RNA accumulates in nuclear foci, while *α*-sat RNA is primarily diffuse in the nucleus, suggesting that when these two satellite RNAs are expressed in both of these cell types, their transcripts may behave in very different ways within the nuclear environment, irrespective of their site of expression within the genome.

**Figure 2.**
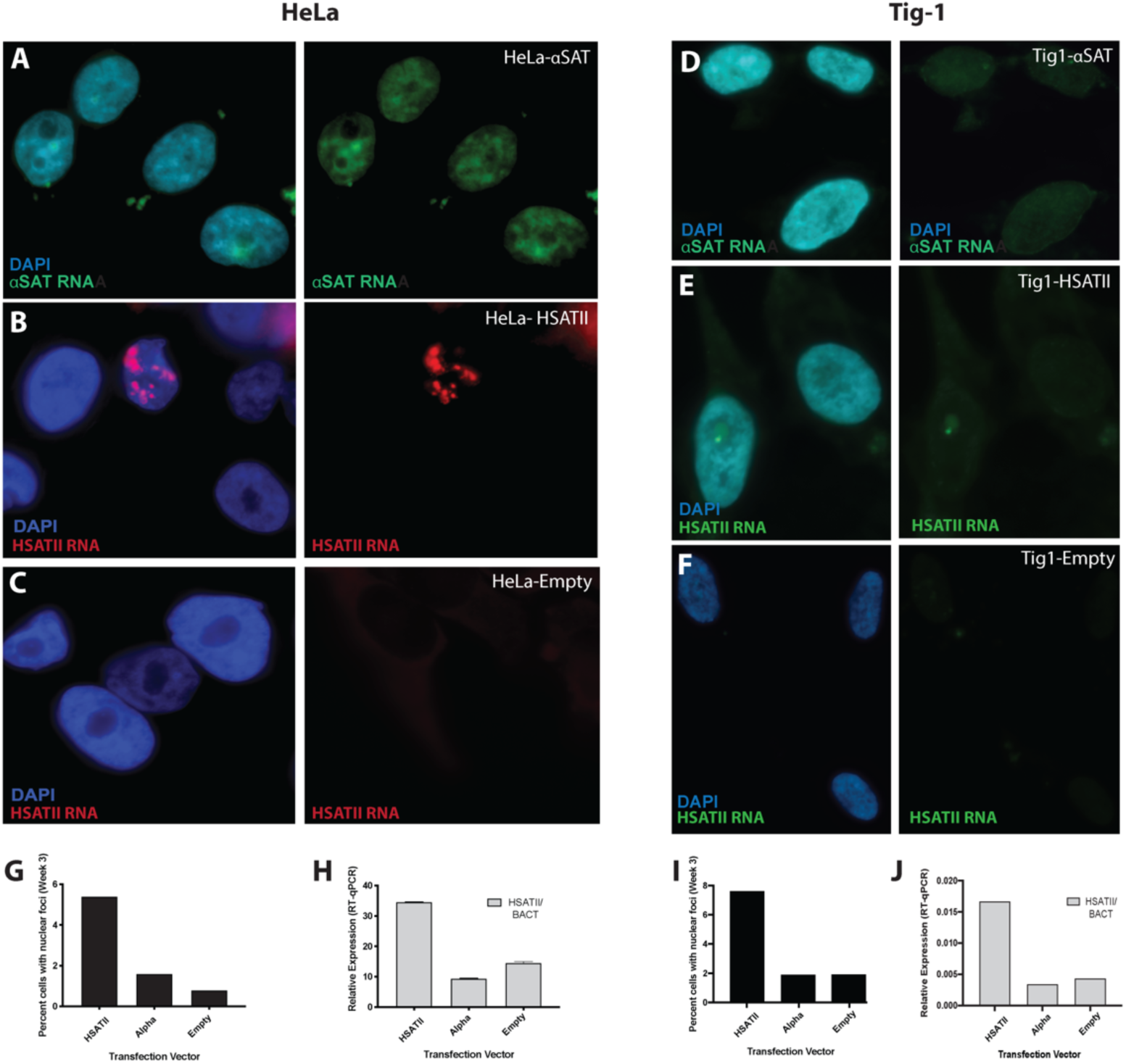
Stable expression of satellite RNA results in focal nuclear accumulation of HSATII RNA in a subset of cells. **A)** Following integration and neomycin selection for the expression construct (HSATII, *α*SAT, empty), a subset of HeLa **(A-C)** and Tig-1 **(D-F)** nuclei exhibit satellite RNA expression by RNA FISH. While *α*-sat RNA **(A, D)** is primarily diffuse in the nucleus, HSATII RNA accumulates in distinct nuclear foci **(B, E)** in both HeLa and Tig-1 cells. Cells transfected with an empty vector have no satellite RNA expression detectable by RNA FISH **(C, F). G, I)**. Quantification of cells with HSATII nuclear expression 3 weeks following transfection in HeLa **(G)** and Tig-1 **(I). (H, J)** RT-qPCR of HSATII RNA in transfected cell lines. Data is shown normalized to ß-actin (housekeeping gene). Error bars show 95% confidence interval.

### HSATII RNA accumulates adjacent to the location from which it is expressed and recruits MeCP2

In cancer cells that express HSATII RNA, the RNA accumulates *in cis* immediately adjacent to their site of transcription (Hall et al., 2017). Therefore, we asked whether the accumulated HSATII RNA foci in stably transfected cell lines also remain *in cis*. Sequential RNA FISH followed by DNA FISH confirmed HSATII RNA accumulated adjacent to the site into which the expression construct (pTargeT backbone) was integrated into the genome in both HeLa (**Figure 3A-B)** and Tig-1 cells **(Figure 3E-F**). While not all sites of integration had a focal accumulation of RNA, when HSATII RNA was detected, it was nearly always adjacent to the location in which the vector had integrated. It is likely that there are genomic locations into which the expression vector has integrated that are not permissible to transcription, that will not tolerate accumulation of the RNA, or for which the CMV promoter may no longer be active. It is also possible that accumulation of HSATII RNA occurs only when proteins are recruited to the RNA to form CAST bodies (Hall et al., 2017), as it is currently unknown whether the RNA alone results in focal accumulations. We observed that focal accumulation of ectopically expressed HSATII RNA is reminiscent of the focal accumulations in cancer cells (CAST bodies) in both their appearance and in the pattern in which they form adjacent to their site of transcription. Therefore, we sought to examine whether HSATII RNA foci recruited MeCP2, one of the protein components within HSATII CAST bodies in cancer cells (Hall et al., 2017), using co-immunofluorescence/RNA FISH. In HeLa cells, nearly all HSATII RNA focal accumulations had an overlapping focal signal with MeCP2 (**Figure 3C-D**). Colocalization analysis of HSATII RNA and MeCP2 in HeLa cells indicated a Pearson correlation coefficient of r∼1, indicating these signals are nearly perfectly overlapping, despite being detected in different fluorescent channels (**Figure 3B)**. This was in contrast to a lack of colocalization of HSATII RNA and pTargeT backbone DNA in HeLa cells (−0.5< r <0.7) (**Figure 3B)**, as was expected based on their adjacency (**Figure 3A)**. In Tig-1 cells, some colocalization with MeCP2 was observed for a subset of HSATII RNA accumulations (**Figure 3G**). However, not all HSATII RNA accumulations in primary fibroblasts recruited MeCP2, suggesting that the cellular context may influence the potential for recruitment of MeCP2 into CAST bodies (**Supplemental Figure 4A**). It might also be possible that additional proteins are recruited to CAST bodies independently of MeCP2, or that HSATII RNA, alone, has the ability to condense into focal accumulations *in cis*. Additionally, there were many focal accumulations of MeCP2 in Tig-1 cells that did not overlap HSATII RNA accumulations (**Figure 3G)**. These MeCP2 foci were also observed in control cells (**Supplemental Figure 4B**) and likely represent normal nuclear accumulations of MeCP2 that do not reside in CAST bodies. These data suggest that ectopically expressed HSATII RNA accumulates *in cis* and remains in nuclear foci near, though not overlapping, their site of genomic integration. Further, these HSATII RNA foci recruit MeCP2, a protein known to be in CAST bodies in cancer cells. Differential MeCP2 recruitment in HeLa and primary fibroblast cells suggests MeCP2 colocalization may be dependent on both the cellular context and/or chromatin environment throughout the nucleus.

**Figure 3.**
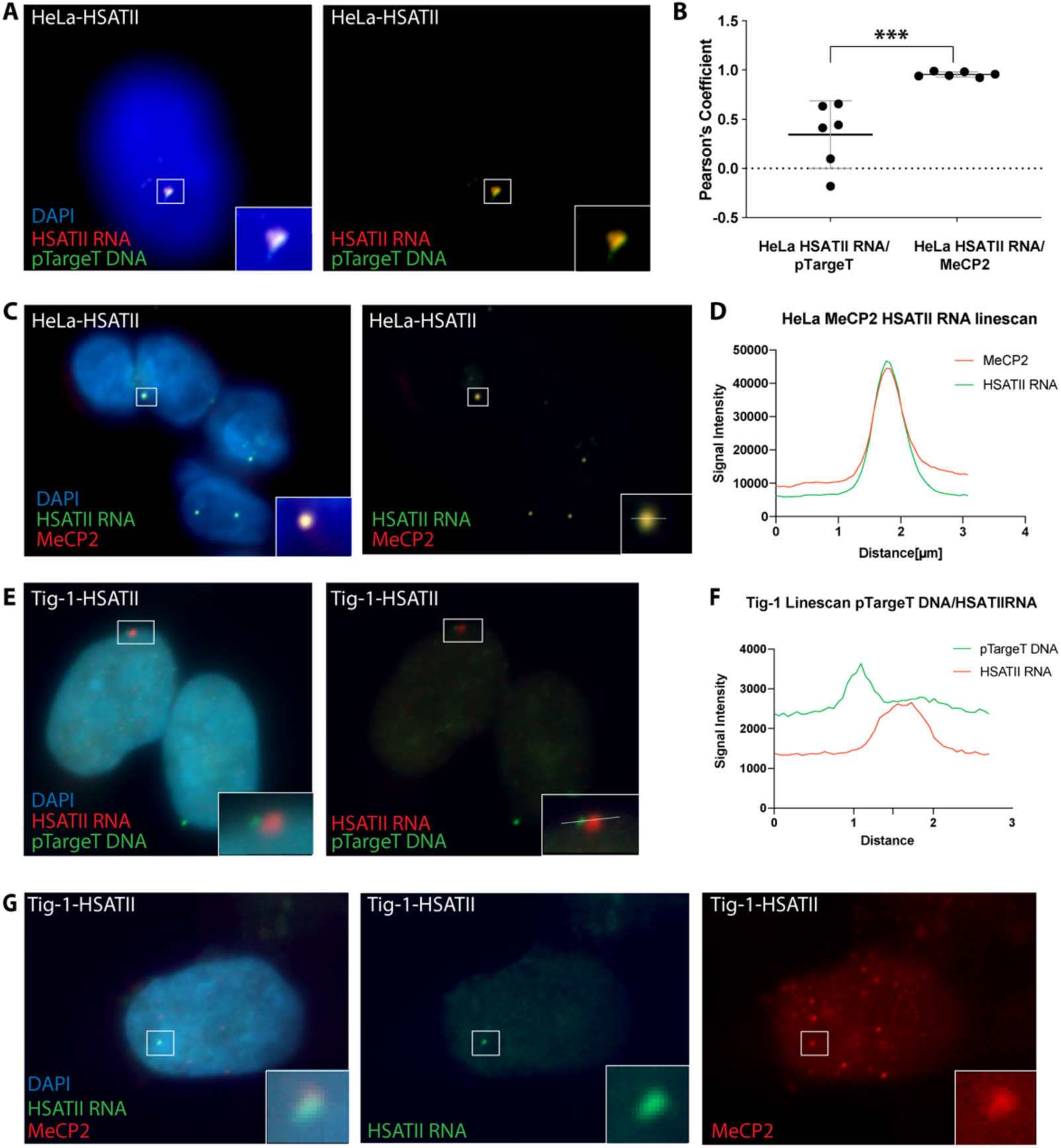
HSATII RNA accumulates adjacent to site of integration and recruits MeCP2. **A)** RNA hyb for HSATII RNA (red) followed by sequential DNA hyb using a probe to detect the pTargeT backbone (green) demonstrates HSATII RNA accumulates primarily adjacent to sites of pTargeT genome integration in HeLa cells stably expressing HSATII 3 weeks after transfection. All boxes with white border indicate regions of interest with zoomed image at bottom right. **B)** Scatter plot of Pearson correlation coefficients for individual nuclei (n=10) hybridized for sequential HSATII RNA/pTargeT DNA and Sat2 RNA with MeCP2 co-IF (Sat2RNA/MeCP2). Black lines indicate mean; grey error bars indicate 95% confidence interval. A t-test indicates a statistical difference (p<.01, n=10) between the correlation coefficients of the two separate analyses (***). **C)** Co-HSATII RNA hyb and IF for MeCP2 indicate a near perfect colocalization of HSATII RNA and MeCp2 signals within HeLa nuclei, as demonstrated by Pearson correlation coefficients plotted in **B. D)** Dual channel linescan of inset shown in **(C)** demonstrating colocalization of MeCp2 (red) and HSATII RNA (green). **E)** RNA hyb for HSATII RNA (red) followed by sequential DNA hyb using a probe to detect the pTargeT backbone (green) demonstrates HSATII RNA accumulates adjacent to, and not overlapping, sites of genome integration in Tig-1 cells stably expressing HSATII 3 weeks after transfection. White line indicates linescan shown in **F)** Dual channel linescan of inset in (**E)** demonstrating the HSATII RNA (red) and pTargeT DNA (green) signals are adjacent, and not overlapping. **G)** Tig-1 fibroblasts stably expressing HSATII RNA for 3 weeks demonstrate overlapping signals for HSATII RNA and MeCP2. Box with white border indicates region of interest with zoomed image at bottom right.

### Satellite RNA expression induces cell division defects and instability

Analysis of cell lines established to stably express HSATII RNA suggests that ectopically expressed HSATII RNA accumulates in a similar manner to HSATII RNA that is endogenously expressed from pericentric regions. It has been demonstrated that aberrant *α*-satellite expression can lead to cell division defects and chromosomal instability (Chan et al., 2017; Ichida et al., 2018; Zhu et al., 2018; Zhu et al., 2011). Therefore, we sought to examine whether cell division defects could be observed in cells that were also forced to express pericentric HSATII, in direct comparison to cells expressing ectopic *α*-sat RNA. Stable cell lines expressing HSATII and *α*-sat RNA were compared to cells with stable integration of the pTargeT backbone alone (empty vector) and control (untransfected) cells at identical passage numbers. Two weeks following transfection, some cell division defects were observed in a small percentage of cells, however this was not significantly different from control cells. In contrast, at 21-28 days following transfection, an increase in chromatin and cell division defects was observed, including an increase in the presence of chromatin bridges (CB), nuclear blebbing/micronuclei (MN), lagging chromosomes (LC), and abnormally shaped nuclei (**Figure 4A**) in HeLa cells. While a significant increase in chromatin bridges was observed in both HSATII and *α*-sat expressing cell lines, an increase in blebbing and overall chromatin defects (sum of all defects observed) was significant (p<.01) only in HSATII-expressing HeLa cells. HeLa are cancer cells, and thus the overall presence of cell division defects in this cell line is predicted to be higher than primary cells. As expected, the frequency of total cell division defects in Tig-1 was much lower than in HeLa (**Figure 4B)**. While the total number of observed cell division defects was not significantly different for either HSATII or *α*-sat expressing cell lines compared to controls, stable Tig-1 cell lines expressing both HSATII and *α*-sat RNA 24 days after transfection exhibited a significant (p<.05) increase in chromatin bridges and nuclear blebbing/micronuclei, which are only very rarely seen (1 cell per 500 cells) in this primary fibroblast cell line when normally maintained at low passage numbers. The increase in cell division and chromatin defects were not simply a result of lipid-mediated transfection since empty vector stable cell lines exhibited a similar number of cell defects as control (untransfected) cells. Primary fibroblast cells exhibited increased cell division defects irrespective of which satellite RNA (HSATII or *α*-sat) was expressed **(Figure 4B**), whereas HeLa cells only exhibited a significant increase in blebbing/MN and abnormally shaped nuclei in HSATII-expressing cells, suggesting that there is an HSATII-expression dependent effect in HeLa cells that is not present in primary fibroblasts (**Figure 4**). This is intriguing in light of previous evidence that HeLa cells exhibit high levels of *α*-sat expression when normally maintained in culture (Hall et al., 2017). Thus, it is possible that HeLa cells respond more marginally to increased *α*-sat expression, whereas the presence of HSATII RNA induces a more drastic range of phenotypes. In contrast, untransfected Tig-1 cells exhibit less *α*-sat expression (Hall et al., 2017), thus may be more sensitive to expression of alpha satellite in the nucleus. Forced alpha satellite expression has been previously shown to induce cell division defects and generate chromosomal instability (Chan et al., 2017; Ichida et al., 2018; Zhu et al., 2018; Zhu et al., 2011), therefore similar mechanisms may be induced upon *α*-sat expression in primary fibroblasts. In HeLa cells, HSATII expression induced a high frequency of blebbing with some resulting in the formation of micronuclei (MN) (**Figure 4A)**, which are a characteristic of cancer cells, but occur much less frequently in primary fibroblasts. Thus, it was surprising that there was a significantly higher incidence of blebbing and micronuclei formation in both HSATII and *α*-sat transfected Tig-1 cells (**Figure 4B**). Further investigation will be required to determine the mechanism by which satellite expression impacts cell division and results in the formation of chromatin bridges and micronuclei, but the observed increase in these cell division aberrations in cells stably expressing *α*-sat or HSATII suggest that expression of either pericentric or centromeric satellite sequences induces cell division defects, and thus may increase chromosomal instability in cancer cells.

**Figure 4.**
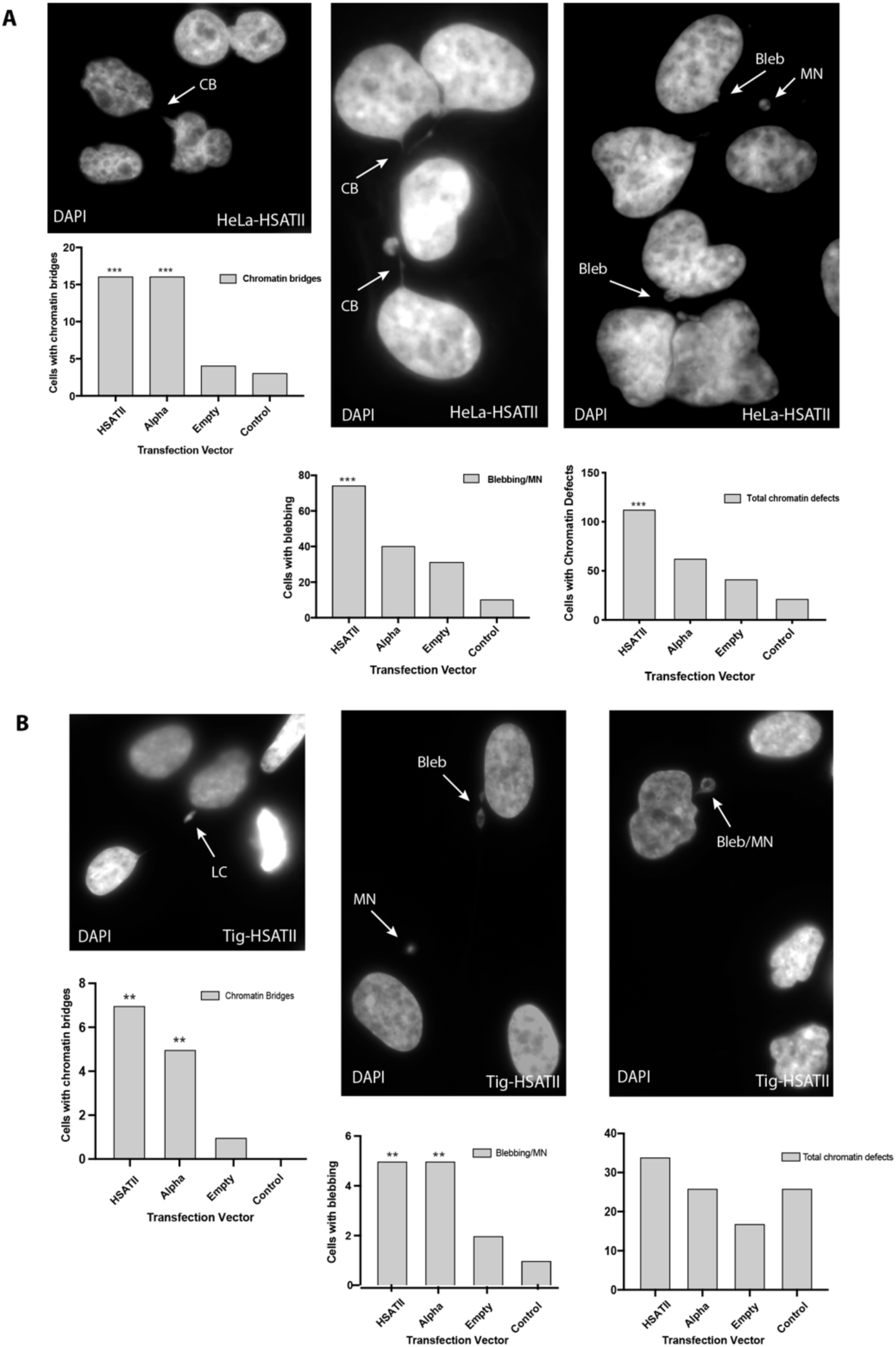
Ectopic expression of satellite RNA leads to cell division defects. **A)** Representative images of HeLa cells transfected and stably expressing HSATII RNA for 3 weeks exhibiting cell division defects including chromatin bridges (CB), blebbing (Bleb) and the formation of micronuclei (MN), as demonstrated in the images in the top 3 panels (DAPI stained, greyscale). Arrows indicate the type of defect observed. Graphs depict counts of cells displaying these defects: chromatin bridges (bottom left), blebbing/MN (bottom middle), and total chromatin defects (bottom right) out of 500 total cells for each transfection vector used. **B)** Representative images of Tig-1 normal fibroblast cells transfected and stably expressing HSATII RNA for 3 weeks displaying cell division defects (lagging chromosomes (LC), blebbing, MN) as indicated by arrows. Graphs depict counts of cells displaying these defects: chromatin bridges (bottom left), blebbing/MN (bottom middle), and total chromatin defects (bottom right) out of 500 total cells for each transfection vector used (X axis). Frequencies of chromatin defects significantly (p<.05) different from control cells are indicated with two asterisks (**). Frequencies of chromatin defects significantly (p<.01) different from control cells are indicated with three asterisks (***).

## Discussion

In order to determine the consequences of the presence of HSATII RNA within cells, we developed a cell culture system to test the effect of HSATII expression by establishing independent cell lines that stably express either HSATII or *α*-sat satellite sequences from randomly integrated sites in the genome. We found that centromeric *α*-sat and pericentromeric HSATII transcripts behave differently when ectopically expressed, yet the presence of HSATII and *α*-sat transcripts both trigger cell division defects. HSATII RNA accumulates in nuclear bodies *in cis*, immediately adjacent to the integration site (**Figure 5**), while *α*-sat RNA is overall more diffuse and does not accumulate appreciably in nuclear bodies, suggesting these two distinct tandemly repeated RNAs have different dynamics within the nuclear environment, and supporting that they may have distinct functions. Nuclear accumulations of HSATII RNA recruit MeCP2 into these nuclear bodies, which are reminiscent of CAST bodies in cancer cells (Hall et al., 2017) (**Figure 5**). Although the transcripts localize in very different ways, expression of both *α*-sat and HSATII induce cell division defects including chromatin bridges, blebbing and micronuclei formation, thus have the potential to generate chromosomal instability and to impact cell division.

**Figure 5.**
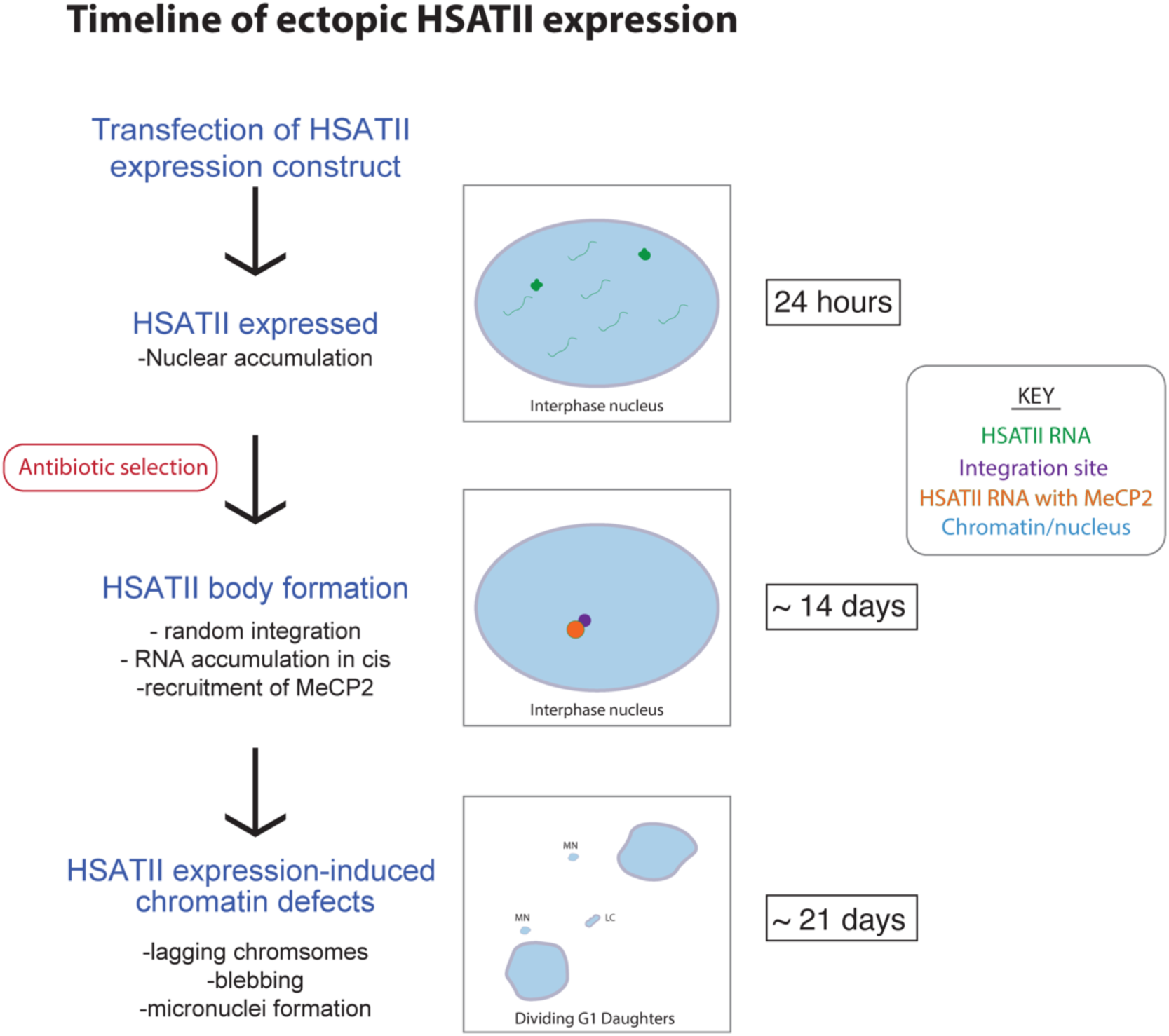
HSATII RNA (green) is expressed transiently 24 hours after transfection in multiple nuclear accumulations and some diffuse nuclear RNA. Following antibiotic selection for 2 weeks (14 days), HSATII RNA bodies form, which recruit MeCP2 (orange), and accumulate adjacent to the genomic integration site of the HSATII expression vector (purple). At this point, HSATII RNA is restricted to these nuclear bodies and little diffuse nuclear RNA is observed. Approximately 3 weeks after transfection (21 days), significant cell division defects are observed, including lagging chromosomes (LC) between dividing daughter cells and the formation of micronuclei (MN).

Previous research has focused on the effect of expression of alpha satellite RNA, which is normally expressed at low levels in all cell types and thought to be involved in centromere protein recruitment (Hall et al., 2017; Johnson et al., 2017; McNulty et al., 2017; Wong et al., 2007). Though a low level of *α*-sat expression is normal and likely necessary for centromere protein recruitment, several groups have described that when *α*-sat is overexpressed, either transiently or by injection of *α*-sat transcripts, this has drastic effects on cell division and leads to chromosomal instability (Chan et al., 2017; Ichida et al., 2018; Zhu et al., 2018), suggesting that the levels of *α*-sat expression are critical to maintaining proper cell division. In contrast, HSATII RNA is not expressed in normal cells, but is highly overexpressed in cancer cells and tissues (Hall et al., 2017; Ting et al., 2011). HSATII RNA accumulates in large nuclear foci within these cells, where it recruits nuclear regulatory proteins including MeCP2. Thus, previous evidence suggested that *α*-sat and HSATII satellite sequences are likely to have unique regulatory mechanisms governing their expression. Our results here suggest that in addition to distinct transcriptional regulatory mechanisms, the transcripts themselves behave in unique ways within the nuclear environment. When an expression vector containing HSATII is randomly integrated into chromosomes, HSATII RNA accumulates immediately adjacent to its integration site and does not appear to drift considerably within the nucleus. Furthermore, these nuclear accumulations recruit MeCP2, which have been previously shown to bind HSATII RNA (Hall et al., 2017). The accumulation of HSATII RNA and recruitment of proteins is reminiscent of “toxic repeat RNAs”, which function to sequester nuclear regulatory proteins in diseases such as frontotemporal dementia/ALS and myotonic dystrophy (Swinnen et al., 2020). Further work will be required to understand the molecular interactions between HSATII RNA and MeCP2, but our results suggest the recruitment of MeCP2 is likely to be a sequence-dependent mechanism, rather than a location-dependent mechanism since HSATII is expressed here from random integration sites in both HeLa and Tig-1 primary fibroblasts (**Figure S2, S3)**.

HSATIII RNA, which is transcribed from Chr 9q12 during stress, assembles into nuclear stress bodies (nSBs), recruiting HSF1 and numerous splicing factors to promote intron retention and suppress splicing of mRNAs during recovery from heat shock (Biamonti and Vourc’h, 2010; Jolly et al., 2004; Ninomiya et al., 2020). Intriguingly, HSATII and HSATIII both derive from CATTC pentamer repeats, despite their divergence and distinct chromosomal localizations. HSATII displays a widespread distribution within the pericentric regions of ∼11 human chromosomes while the bulk of HSATIII resides on Chr 9 and Y, with some smaller loci interspersed within pericentric regions of additional chromosomes (Altemose et al., 2014; Tagarro et al., 1994). HSATII and HSATIII RNA appear to recruit different protein binding partners, yet they both display conserved functional aspects in their ability to accumulate *in cis* and recruit/sequester protein binding partners. It is possible that the tandemly repetitive nature of pentamers embedded within HSATII/HSATIII repeats may facilitate this recruitment given that identical sequences (monomers) are present in high copy number within a linear array. It is possible to envision that if a sequence binding motif is resident within one monomer, that same sequence is also in neighboring monomers, and that sequence would have equal capacity to bind that same protein. Thus, the potential for tandemly repeated sequences to “soak up” protein binding partners exists. In normal cells, HSATII is resident with large blocks of heterochromatin and no HSATII RNA is transcribed. In cancer cells, when pericentric heterochromatin structure is compromised by loss of DNA methylation (Ehrlich, 2009; Hall et al., 2017) and other heterochromatin marks (Slee et al., 2012; Tasselli et al., 2016), there then exists the possibility for expression, and for the resulting RNA to act as a molecular sponge. Our results here suggest this ability to accumulate and sequester regulatory proteins may be a sequence or length-dependent mechanism, since *α*-sat, which is similar in its tandemly repeated nature, yet of a 171-bp monomer length not consisting of CATTC repeats, does not appear to accumulate or recruit proteins upon random integration and ectopic expression.

Nuclear stress bodies form on HSATIII noncoding RNA and recruit proteins into nuclear condensates, a theme which is emerging as a conserved feature of RNP granules that are likely to exist within liquid-liquid phase separated domains in the nucleus. Recent work highlights the role of RNA in modulating the biophysical properties of liquid droplets, and the accumulation of misregulated transcripts may lead to mislocalization of RNA binding proteins containing intrinsically disordered domains (IDRs), such as FUS and TDP-43 (Maharana et al., 2018). Further, RNA itself may facilitate the nucleation of these liquid droplets, in a size and concentration-dependent manner (Garcia-Jove Navarro et al., 2019). This intrinsic property of RNA may help to explain why lncRNAs, such as Neat-1, can nucleate phase separation into paraspeckles upon interaction with specific proteins containing IDRs (Hennig et al., 2015). MeCP2, though classically described as a DNA binding protein, has been more recently identified in an RNA-binding protein (RBP) screen (Castello et al., 2016) and binds mRNAs in the brain, where it affects alternative splicing (Young et al., 2005). MeCP2 also contains an IDR, and appears to bind and colocalize with HSATII RNA within CAST bodies, which we note also resemble nuclear condensates. MeCP2 and its binding partner, Sin3A, were previously observed to bind HSATII, but additional work will be required to identify if other proteins are also recruited to CAST bodies, and if these HSATII RNA-nucleated bodies form liquid-liquid phase separated domains within the nucleus.

Since initially found to be expressed in cancer cells, the functional consequence of HSATII transcripts within the nucleus has remained elusive, however induction and injection of HSATII RNA was found to result in the formation of cDNA intermediates, which can facilitate the expansion of pericentromeric HSATII-containing DNA at endogenous sites in the genome (Bersani et al., 2015). Though the mechanism facilitating this copy number gain is not well understood, it is thought to result from the formation of RNA-DNA hybrids following reverse transcription of HSATII RNA transcripts. In addition to cancer cells, HSATII RNA has recently been found to be induced upon herpesvirus infection, where it is thought to be involved in the viral life cycle (Nogalski et al., 2019) and in human FSHD cell models where it resides in nuclear dsRNA foci that also aggregate proteins (Shadle et al., 2020). HSATII RNA has also been implicated in the growth of cancer cells (Ting et al., 2011), and its expression is further induced in cells grown under 3D culturing conditions (Bersani et al., 2015). Thus, there may be convergent mechanisms by which HSATII RNA can more globally affect both growth and viral replication. Though these effects have been observed, the specific role that HSATII transcripts play in mediating these phenotypes has remained unclear. Our observation that HSATII RNA accumulates and recruits regulatory protein *in cis*, irrespective of the site from which it is expressed, may shed some significant insight into the molecular function of HSATII RNA. The ability of HSATII RNA to bind, and potentially sequester, large amounts of regulatory proteins is likely to have a large effect on transcription and genome regulation more broadly, and may ultimately lead to widespread heterochromatin instability, which is observed in cancer and other diseases (Carone and Lawrence, 2013).

Supporting a more global effect of HSATII accumulations within the nucleus, our results implicate HSATII transcripts in generating cell division defects, many of which are also known to be of higher frequency in cancer cells. When HSATII RNA is expressed ectopically in both HeLa and Tig-1 primary fibroblasts for several weeks, we observed increased frequencies of chromatin bridges, lagging chromosomes, and micronuclei formation (**Figure 4**). These phenotypes have previously been observed upon introduction of satellite transcripts into both mouse and human cells, where they may induce DNA damage via the formation of RNA-DNA hybrids (Zhu et al., 2018; Zhu et al., 2011). Overexpression of *α*-sat has been linked to chromosomal instability and segregation errors resulting in copy number changes in daughter cells (Ichida et al., 2018). Our results here support previous links between centromeric satellite expression and chromosomal instability (Bouzinba-Segard et al., 2006; Chan et al., 2017; Ichida et al., 2018; Slee et al., 2012), and extend this phenomenon to expression of pericentromeric HSATII RNA.

Despite their abundant presence within pericentromeres, the role of tandemly repeated satellite sequences within these regions has been elusive. Recent work suggests pericentric satellites within fruit fly and mouse heterologous chromosomes may act to tether chromosomes within nuclei via the formation of chromocenters (Jagannathan et al., 2018). Perturbation of proteins within chromocenters results in the formation of micronuclei due to budding from the nucleus, supporting that tethering of pericentric regions to chromocenters is key to nuclear organization and overall nuclear integrity (Jagannathan et al., 2018). Alteration of epigenetic regulatory mechanisms within heterochromatin is also known to cause chromosomal instability (Slee et al., 2012), thus impaired maintenance of heterochromatin is likely to lead to both satellite expression and chromosomal instability. Our results here add to a growing body of evidence suggesting that the presence of satellite transcripts, themselves, are likely to induce defects in chromosome segregation. This observation is supported by previous studies of transient *α*-sat expression, but the more long-term and specific effect of the presence of satellite RNA has been unclear. By randomly integrating and expressing ectopic HSATII and *α*-sat RNA, we demonstrate the ability to observe the specific behavior of these satellite transcripts within the nucleus, irrespective of their site of transcription. Despite displaying strikingly different nuclear localization, the presence of both *α*-sat and HSATII RNA leads to chromosome mis-segregation and the formation of micronuclei, suggesting convergent roles for these transcripts in the perturbation of cell division.

## Materials and Methods

**Key Resource Table**

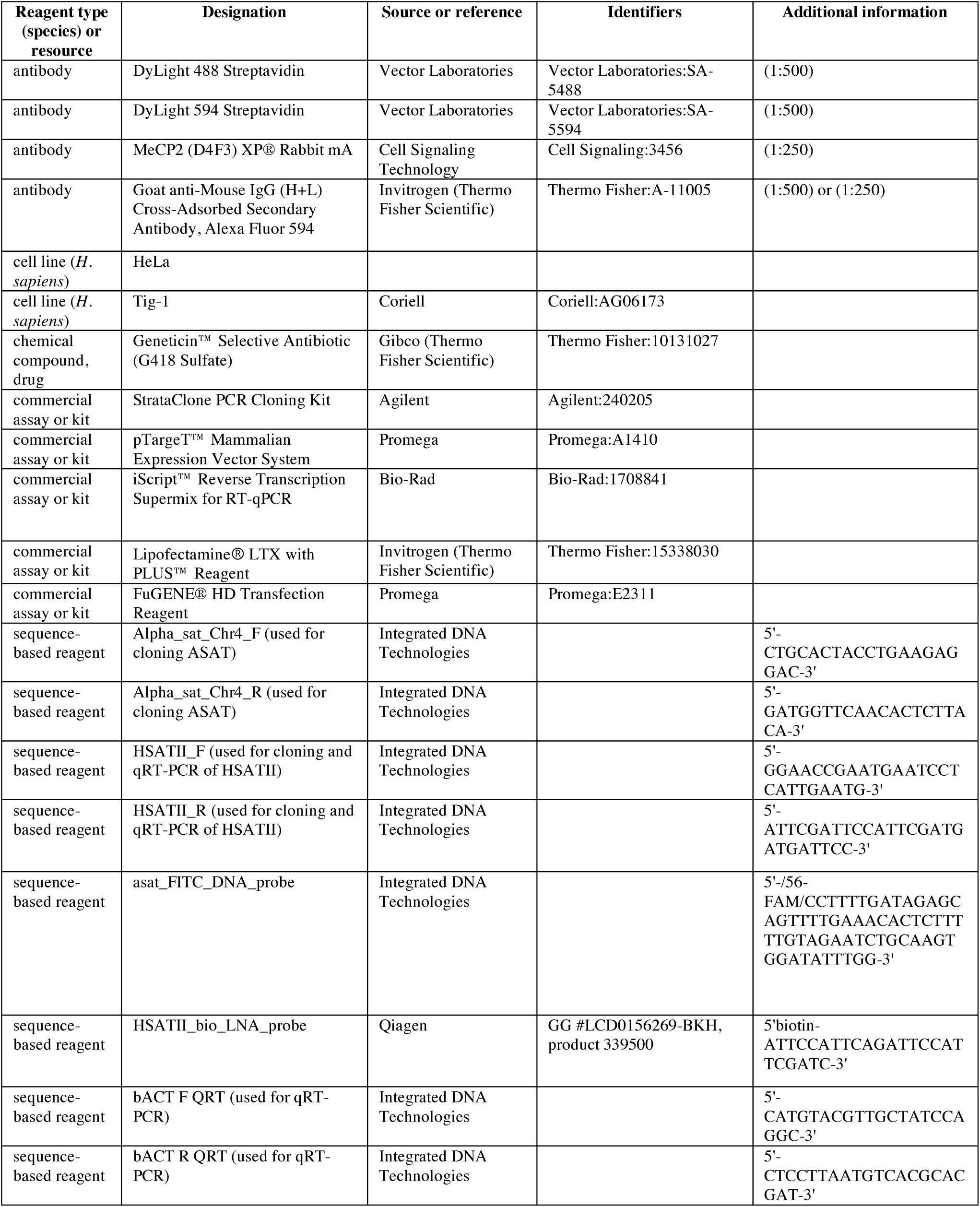
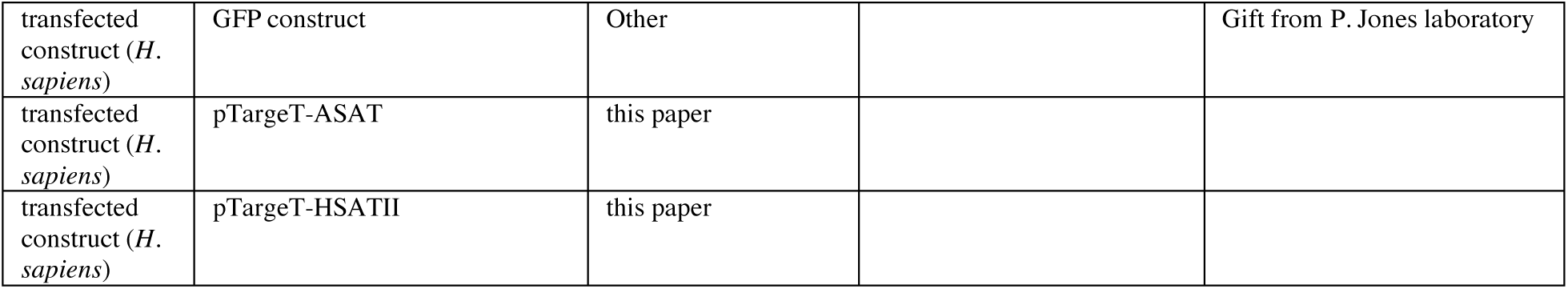

### Construction of DNA plasmids

The pTargeT™ Mammalian Expression Vector System (Promega, Madison, WI), containing a CMV promoter and neomycin resistance, was used for transfection and long-term selection of stably transfected lines. Insert DNA was derived from total RNA extracted from U2OS cells and then reverse transcribed using an iScript™ Reverse Transcription Supermix (Bio-Rad Laboratories, Hercules, CA). A 140 bp region of alpha satellite sequence from chromosome 4 and a 349 bp region of HSATII from chromosome 7 were independently cloned into pTargeT™ DNA backbone via StrataClone TA cloning (Agilent Technologies, Santa Clara, CA).

### Cell culture and transfection

HeLa human epithelioid cervix carcinoma cells and Tig-1 human primary fibroblast cells (Coriell AG06173) were cultured in the presence of 5% CO_2_ prior to transfection in Minimal Essential Medium (Gibco, Thermo Fisher Scientific, Waltham, MA) supplemented with 10% (HeLa) or 15% (Tig-1) fetal bovine serum (HyClone, GE Healthcare Life Sciences, Marlborough, MA; Avantor Seradigm, VWR, Radnor, PA), 100 units/mL penicillin and 100 µg/mL streptomycin (Gibco, Thermo Fisher Scientific) and 2 mM L-glutamine (Gibco, Thermo Fisher Scientific). After transfection, cells were maintained in the above media, replacing the penicillin/streptomycin with geneticin (G418 sulfate) (Gibco, Thermo Fisher Scientific) at a concentration of 700 µg/mL for HeLa cells and 500 µg/mL for Tig-1 cells for selection over a minimum of 14 days.

Transient transfections of the empty pTargeT™ vector, pTargeT-*α*SAT and pTarget-HSATII were performed in parallel in HeLa cells using Lipofectamine^®^ LTX with PLUS™ Reagent (Invitrogen, Thermo Fisher Scientific). Cells were seeded on 22×22mm coverslips or in T-25 flasks such that they were 70-90% confluent at the time of transfection. The plasmid DNA-lipid complexes for transfection were prepared following the manufacturer’s protocol. For cells in 6-well plates, 2.5 µg DNA was transfected per well using a 2.5µg:22.5 µL ratio of plasmid DNA:Lipofectamine^®^ LTX, supplemented with 3.5µL PLUS™ Reagent. Cells in T-25 flasks were transfected with 3 µg plasmid DNA per flask, with 312.5 µL Lipofectamine^®^ LTX and supplemented with 12.5 µL PLUS™ Reagent. 24 hours following transfection, cells on coverslips were fixed for FISH experiments and cells in flasks were pelleted and frozen for RNA extraction and qRT-PCR analysis.

Stably transfected lines of HeLa cells were cultured in T-25 flasks, also using a ratio of 3 µg plasmid DNA:315 µL Lipofectamine^®^ LTX and 12.5 µL PLUS™ Reagent per flask. In parallel with the flasks transfected with the empty pTargeT™ vector, pTargeT-*α*SAT and pTargeT-HSATII plasmids, untransfected HeLa cells was cultured in parallel as a control. Cells were transfected 24 hours after seeding in flasks, and then maintained in standard growth medium containing 10% FBS for 3 days before adding G418 sulfate-supplemented media. Cells were cultured in selective media for 2 weeks, during which the media was exchanged every 2 days. After this selection period, the stably transfected lines were maintained for an additional several weeks. Cells were then plated on coverslips and fixed for FISH experiments twice a week, and harvested 3 and 4 weeks following the end of selection, for RNA extraction and qRT-PCR analysis.

To optimize lipid-mediated transfection for Tig-1 primary fibroblasts for both expression levels as well as overall health of transfected cells, a GFP expression vector (obtained from P. Jones) was transiently transfected into cells on coverslips using a range of conditions, with 3 µg plasmid DNA transfected per well of a 6-well plate. Lipofectamine^®^ LTX was tested at volumes of 9 µL and 12 µL per well, and FuGENE® HD was tested at a range of concentrations (3:1, 4:1 and 5:1 per μg DNA). After 24 hours, media was exchanged and cells maintained in standard media for an additional 24 hours. Cells were then visualized in culture prior to fixation and DAPI staining as described below. Transfection efficiency was evaluated by scoring GFP expression and cellular health gauged by scoring the number of cells per field of view.

Tig-1 cells were stably transfected using FuGENE® HD Transfection Reagent (Promega), following the manufacturer’s protocol. Cells in T-25 flasks were grown to 60% confluency and then transfected using 8 µg plasmid DNA and 44 µL FuGENE® HD Transfection Reagent per flask. Three control flasks were cultured in addition to the pTargeT lines: a wild type untransfected flask not treated with selective media, a wild type untransfected flask selected with G418 sulfate, and a mock flask treated with the transfection reagent and then selected with G418 sulfate. Media containing transfection reagent was removed 24 hours post transfection and replaced with selective media. Selection was complete after 10-14 days, after which the stably transfected cells were maintained for several weeks. During this time, cells were fixed on coverslips weekly and harvested for RNA extraction at multiple time points.

### Preparation of chromosome spreads

Chromosome spreads were prepared as previously described (Howe et al., 2014), with the following modifications. Cells in flasks were treated with colcemid for 1-2 hours depending on the cell line and monitored for mitotic cells, at which point the cells were harvested, treated with hypotonic solution (0.075M KCl) and fixed with Carnoy’s Fixative (3:1 ratio of methanol:glacial acetic acid). Fixed cells were then dropped onto clean glass slides under humid conditions to produce chromosome spreads. Slides were serially dehydrated with ethanol and stored at -20°C. Thawed slides were dehydrated in 100% ethanol prior to DNA FISH experiments.

### Fluorescence *in situ* hybridization (FISH) and immunostaining

For all FISH and immunostaining on interphase cells, transfected cell lines were grown on 22×22 mm coverslips and then fixed as follows: 3-5 minutes in cytoskeletal buffer (Byron et al., 2013) plus 0.5% Triton X-100 (Sigma-Aldrich, St. Louis, MO) and 10 mM ribonucleoside vanadyl complex (New England Biolabs, Ipswich, MA), 10 minutes in 4% paraformaldehyde (Ted Pella, Redding, CA) in 1x PBS, followed by storage in 70% ethanol at 4°C.

RNA FISH was performed using a FITC-labeled DNA oligo probe complementary to alpha satellite (Integrated DNA Technologies, Newark, NJ) or a biotinylated locked nucleic acid (LNA) oligo probe complementary to HSATII (QIAGEN, Hilden, Germany) (see Key Resources Table). For each coverslip, 0.25 pmol of oligonucleotide was denatured for 10 minutes at 80°C in 30% formamide, then diluted in hybridization buffer to a final concentration of 15% formamide. Hybridization buffer contained 2 mg/mL bovine serum albumin (BSA) (Roche, Basel, Switzerland), 10% dextran sulfate and 2x SSC, supplemented with RNasin Plus RNase Inhibitor (Promega). The probe was then applied to coverslips, incubated overnight in a humid chamber at 37°C and then washed for 15 minutes in 15% formamide in 2x SSC at 37°C, 15 minutes in 2x SSC at 37°C, 15 minutes in 1x SSC at RT, and 5 minutes in 4x SSC at RT. HSATII RNA was detected with either 1:500 DyLight 488 streptavidin or DyLight 594 streptavidin (Vector Laboratories, Burlingame, CA) in 4x SSC+1% BSA. Secondary-detected coverslips were then washed at room temperature for10 minutes in 4x SSC, 10 minutes in 4x SSC+0.1% Triton X-100, and 10 minutes in 4x SSC. All coverslips were stained with 2 µg/mL DAPI, mounted with VECTASHIELD® Antifade Mounting Medium (Vector Laboratories), and sealed with fingernail polish.

A pTargeT™ vector was labeled by nick translation with digoxigenin-11-dUTP (Roche) (Byron et al., 2013) to detect integration of the construct via DNA FISH in both interphase cells and mitotic chromosome spreads. To prepare the probe for hybridization, 50 ng of nick translated probe per coverslip was combined with 12 μg of human Cot-1 DNA (Roche), 10 μg salmon sperm ssDNA (Sigma-Aldrich) and 20 μg *E. coli* tRNA (Sigma-Aldrich), dried with a speed-vac, and then resuspended in 10 μL formamide. Coverslips were first treated with 0.2 N NaOH in 70% ethanol to remove RNA, then dehydrated in 100% ethanol and air dried. Interphase cells or chromosome spreads were denatured in 70% formamide in 2x SSC, pH 7.0, for 2 minutes at 80°C, followed by 5 minutes in cold (4°C) 70% ethanol and 5 minutes in cold 100% ethanol prior to air drying. Probes were diluted in the hybridization buffer used for RNA FISH, to a final concentration of 50% formamide, before applying to coverslips and incubating overnight at 37°C in a humid chamber. Coverslips were then washed as for RNA FISH, with the adjustment of the first wash to 50% formamide/2x SSC. The probe was detected with anti-digoxigenin-fluorescein (Roche), diluted 1:500 in 4x SSC+1% BSA. Secondary incubation, washes, DAPI staining and mounting were carried out as for the RNA oligo hybridization. RNA and DNA co-hybridization was performed by first completing the RNA hybridization as described, followed by fixation in 4% paraformaldehyde for 10 minutes prior to DNA FISH, with the removal of the NaOH treatment step.

In HeLa cells, MeCP2 and HSATII RNA co-visualization was performed by first hybridizing oligo probes to RNA as described, followed by a 10-minute fixation in 4% paraformaldehyde prior to staining for MeCP2. For Tig-1 cells, MeCP2 was stained first, then the signal was fixed in 4% paraformaldehyde for 10 minutes before proceeding with the RNA FISH protocol. The HSATII probe was detected with DyLight 488 streptavidin and the MeCP2 with an Alexa fluor 594 goat anti-rabbit secondary. To stain for MeCP2, coverslips were rinsed for 10 minutes in 1x PBS, then incubated with a 1:250 dilution of MeCP2 rabbit antibody (Cell Signaling Technology, Danvers, MA) in 1x PBS+1% BSA at 37°C for 1-3 hours. Coverslips were washed at room temperature: 10 minutes in 1x PBS, 10 minutes in 1x PBS+0.1% Triton X-100, and 10 minutes in 1x PBS. The secondary antibody (diluted in 1x PBS+1% BSA, (1:500) for HeLa cells, and (1:250) for Tig-1 cells) was then applied and incubated as for the primary, with the same set of washes. Following both MeCP2 staining and HSATII RNA hybridization, coverslips were DAPI stained and mounted.

### Image acquisition and analysis

Slides were imaged using a ZEISS Axio Observer Z1 epifluorescent microscope equipped with a Hamamatsu ORCA-Flash 4.0 Digital CMOS camera or ZEISS Axiocam 702 mono camera with a 100x oil objective. Both single plane images and Z-stacks were captured and analyzed using ZEISS ZEN2 imaging software.

Counts of nuclear body formation or chromatin defects were scored manually, while colocalization was measured using the ZEN2 Colocalization Module, including generation of Pearson’s correlation coefficient. Linescans were generated using the Profile tool in ZEN2. DNA integration foci from pTargeT DNA hybridizations on interphase cells were scored on a subjective scale for relative intensity as determined by observation through the microscope oculars. This scale categorized relative foci brightness into the following five subjective classes: 0 (no foci), + (dim), ++ (easily visible), +++ (very bright), or ++++ (extremely bright). DNA integration foci from chromosome spreads hybridized with the pTargeT probe were categorized as being on acrocentric or submetacentric chromosomes, and further scored based on signal location relative to the centromere as well as chromosome size. Chromatin defects were scored by imaging DAPI stained slides and evaluating nuclei for abnormal phenotypes. Categories for these defects included: chromatin bridges between two nuclei (including lagging chromosomes), micronuclei which were fully separated from the primary nucleus, abnormally shaped nuclei (including nuclei in the process of blebbing) and other defects (including nuclei with burst chromatin and holes or indentations in the chromatin). All scoring data were analyzed and graphed using Prism8 (GraphPad), including statistical analysis.

### Quantitative RT-PCR analysis

RNA was extracted from pelleted cells using either an RNeasy Mini kit and QIAshredder (QIAGEN) or a Direct-zol RNA Miniprep kit (Zymo Research, Irvine, CA), following manufacturers’ protocols for each. Samples were then treated with TURBO™ DNase (Invitrogen, Thermo Fisher Scientific) following manufacturer guidelines. RNA was purified using Agencourt AMPure XP beads (Beckman Coulter, Brea, CA) and quantified via NanoDrop™ (Thermo Fisher Scientific). Reverse transcription was performed using iScript™ Reverse Transcription Supermix for RT-qPCR (Bio-Rad), including RT-controls for all RNA samples. Quantitative RT-PCR was carried out with PerfeCTa® SYBR® Green FastMix® Reaction Mix (Quantabio, Beverley, MA), using primers included in the Key Resources Table. All reactions were performed in triplicate, using a CFX96 Real-Time C1000 Thermal Cycler (Bio-Rad) or MJ Research PTC-200 thermocycler equipped with a Chromo4 Real-Time PCR Detector (Bio-Rad). Analysis of relative enrichment was calculated using the ΔΔC(t) method (Pfaffl, 2001), where reported values are normalized to β-actin expression.

## Acknowledgements

This work was supported by funding from the Charles E. Kaufman Foundation of the Pittsburgh Foundation (KA2017-91790) and the National Institutes of Health (R15 GM134495) to DMC. We would like to thank S. Akkipeddi for assistance with maintenance of stably transfected cell lines and H. Sen for assistance with chromosome spreads and DNA hybridizations. We are grateful to J. Rubien, S. Akkipeddi, and B. Carone for helpful comments and critical reading of the manuscript.

