## Supplemental Figures for "Ectopic expression of pericentric HSATII RNA results in nuclear RNA accumulation, MeCP2 recruitment, and cell division defects"

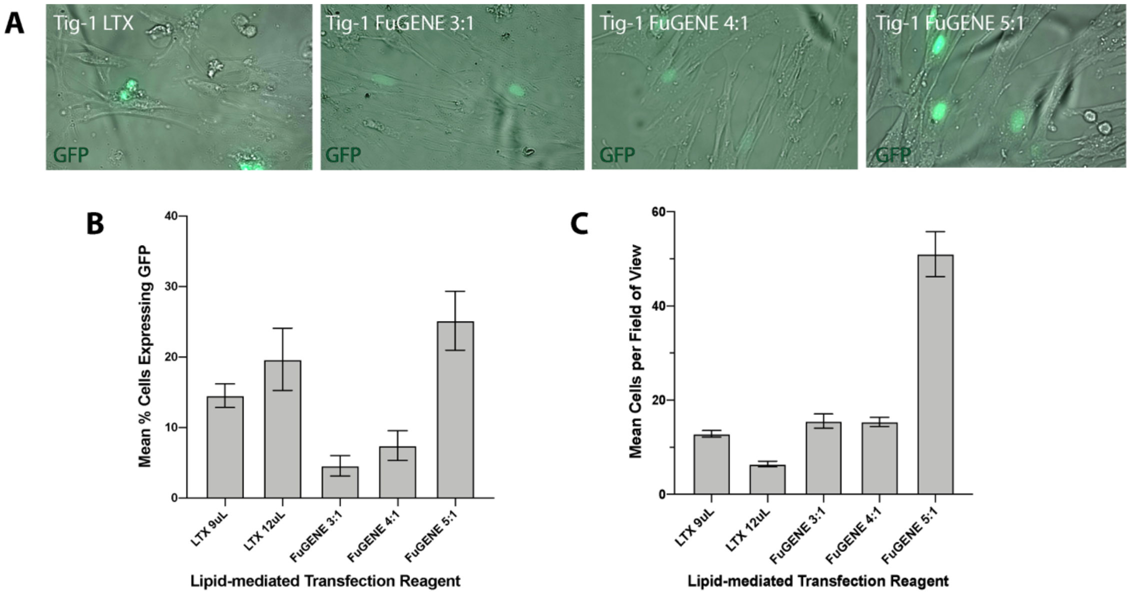


**Figure S1. Optimization of lipid-mediated transfection protocol for primary human fibroblasts. A)** Representative images of transient transfection of Tig-1 cells with GFP expression vector and (left to right): 9 uL of Lipofectamine LTX reagent, 3:1 ratio of FuGENE HD reagent to GFP vector, 4:1 ratio of FuGENE HD, and 5:1 ratio of FuGENE HD reagent to GFP vector. **B)** Transfection efficiencies using Lipofectamine LTX or FuGENE HD reagent in scaled concentrations. Mean percentage of cells with GFP expression out of total cells per field of view (± SEM). **C)** Cell viability of selected reagents and concentrations on transfected Tig-1 cells. Mean number of cells per field of view is shown (± SEM).

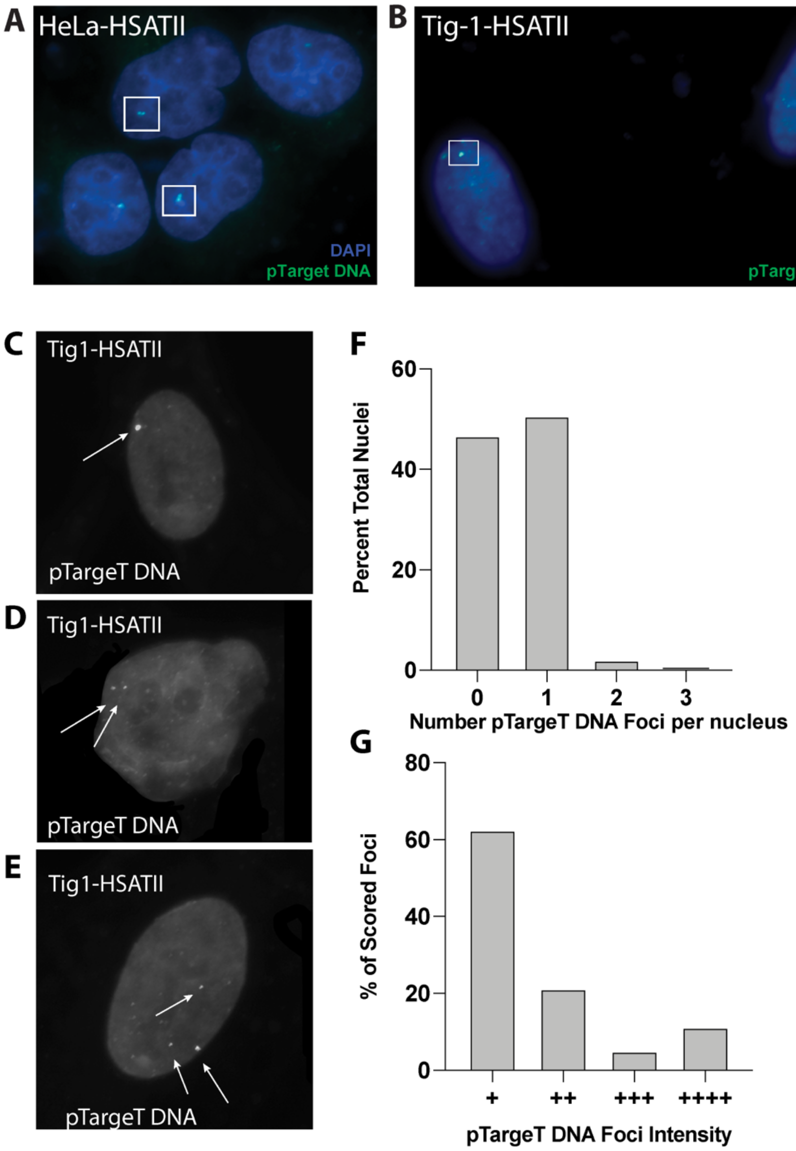


**Figure S2. pTargeT vector DNA integration at high efficiency in HSATII transfected Tig-1 cells. A-B)** Hybridization of pTarget (empty vector) shows integration into one or two genomic loci per nucleus in both HeLa **(A)** and Tig-1 **(B)** nuclei. FISH with labeled pTargeT backbone probe for Tig-1 showing **C)** Representative HSATII transfected nucleus with one site of integration. **D)** Representative HSATII transfected nucleus with two sites of integration**. E)** Representative HSATII transfected nucleus with three sites of integration. **F)** DNA FISH with the pTargeT backbone shows at least one site of pTargeT integration in 54.69% of HSATII transfected cells (N = 200). Due to incomplete DNA hybridization efficiency, not all nuclei displayed detectable pTargeT DNA signal. **G)** Relative intensity of scored foci was assigned to the following categories: + (dim), ++ (easily visible), +++ (very bright), or ++++ (extremely bright, saturated pixels). All foci were categorized and imaged using the same exposure time.

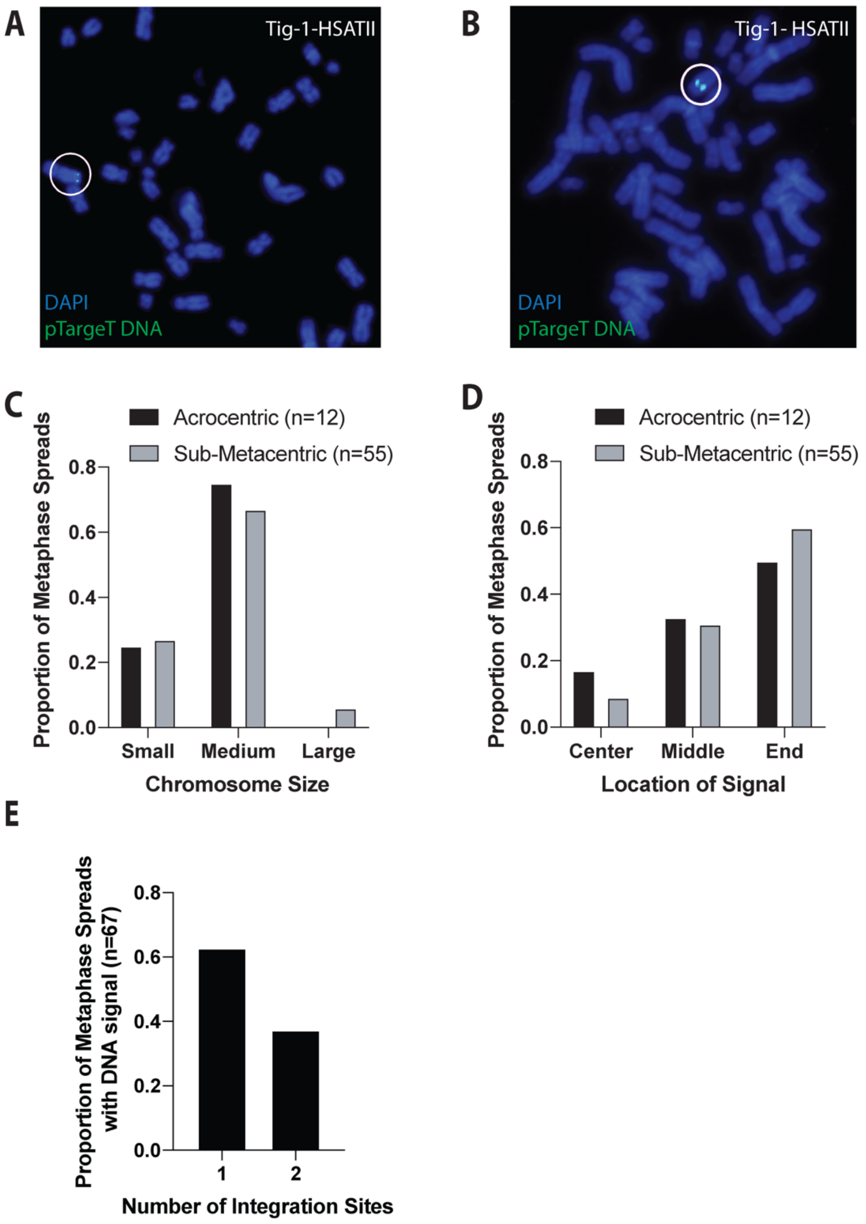


**Figure S3. Stable Tig-1 cell lines display random integration of pTargeT vector DNA into Tig-1 chromosomes. A-B)** Labeled pTargeT backbone probe detects integration sites of transfection vector containing HSATII DNA construct (green) in both **(A)** sub-metacentric and **(B)** acrocentric chromosomes on DAPI-stained metaphase spreads. Integration sites are indicated by circles and have been replicated on sister chromatids. **C-D)** Categorization of **(C)** chromosome size and **(D)** signal location were scored for both sub-metacentric and acrocentric chromosomes and demonstrate random integration on a variety of human chromosomes. Since most human chromosomes are categorized as "medium" size, the higher frequency of integration into medium-sized chromosomes likely reflects a random distribution of integration sites. **E)** The number of pTargeT vector DNA integration sites on HSATII transfected Tig-1 chromosome spreads was also scored. Data shown is for chromosome spreads with detectable pTargeT DNA hybridization signal.

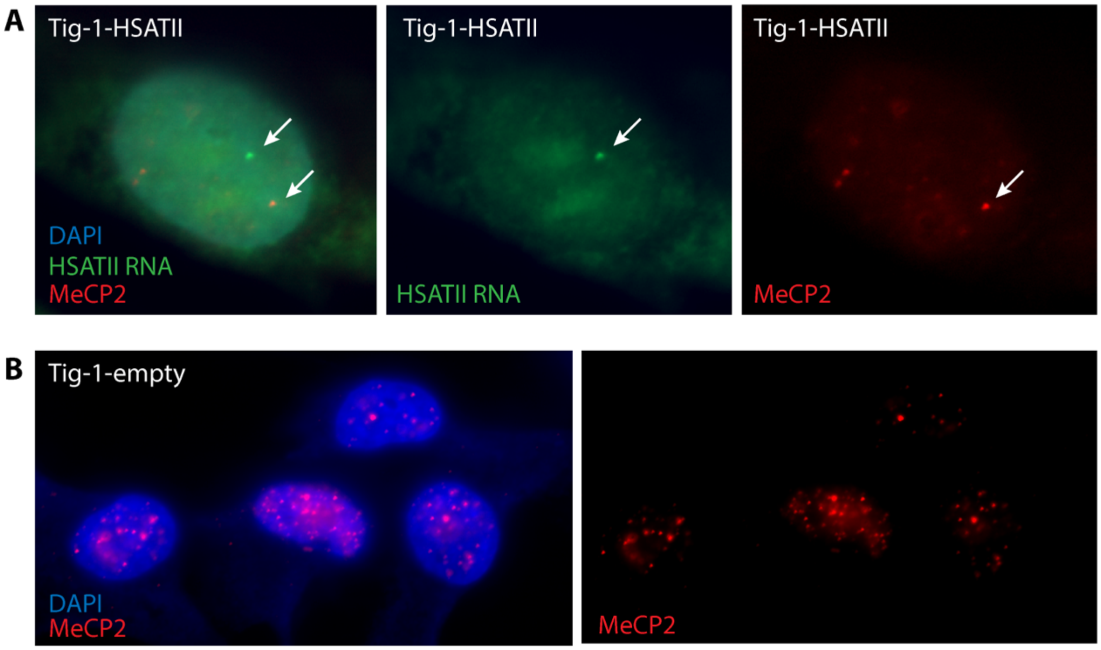


**Figure S4. MeCP2 localizes with a subset of HSATII foci in transfected Tig-1 cells. A)** Tig-1 cells transfected and expressing HSATII (green) do not always recruit MeCP2 (red) to HSATII RNA accumulations (HSATII and MeCP2 distinct foci indicated by separate arrows). **B)** Tig-1 cells transfected with empty vector (pTargeT only) demonstrate MeCP2 nuclear foci (red) in a normal nuclear distribution.
